# A deep learning-based approach for unbiased kinematic analysis in CNS injury

**DOI:** 10.1101/2024.04.08.588606

**Authors:** Maureen Ascona, Ethan Kim Tieu, Erick Gonzalez-Vega, Daniel J Liebl, Roberta Brambilla

## Abstract

Traumatic spinal cord injury (SCI) is a devastating condition that impacts over 300,000 individuals in the US alone. Depending on the severity of the injury, SCI can lead to varying degrees of sensorimotor deficits and paralysis. Despite advances in our understanding of the underlying pathological mechanisms of SCI and the identification of promising molecular targets for repair and functional restoration, few therapies have made it into clinical use. To improve the success rate of clinical translation, more robust, sensitive, and reproducible means of functional assessment are required. The gold standards for the evaluation of locomotion in rodents with SCI are the Basso Beattie Bresnahan (BBB) and Basso Mouse Scale (BMS) tests.

To overcome the shortcomings of current methods, we developed two separate marker-less kinematic analysis paradigms in mice, MotorBox and MotoRater, based on deep-learning algorithms generated with the DeepLabCut open-source toolbox. The MotorBox system uses an originally designed, custom-made chamber, and the MotoRater system was implemented on a commercially available MotoRater device. We validated the MotorBox and MotoRater systems by comparing them with the traditional BMS test and extracted metrics of movement and gait that can provide an accurate and sensitive representation of mouse locomotor function post-injury, while eliminating investigator bias and variability. The integration of MotorBox and/or MotoRater assessments with BMS scoring will provide a much wider range of information on specific aspects of locomotion, ensuring the accuracy, rigor, and reproducibility of behavioral outcomes after SCI.

**Highlights:**

- MotorBox and MotoRater systems are two novel marker-less kinematic analysis paradigms in mice, based on deep-learning algorithms generated with DeepLabCut.
- MotorBox and MotoRater systems are highly sensitive, accurate and unbiased in analyzing locomotor behavior in mice.
- MotorBox and MotoRater systems allow for sensitive detection of SCI-induced changes in movement metrics, including range of motion, gait, coordination, and speed.
- MotorBox and MotoRater systems allow for detection of movement metrics not measurable with the BMS.

## 1. Introduction

Traumatic spinal cord injury (SCI) is a devastating condition that occurs when a mechanical force (primary injury) causes direct damage to the spinal cord tissue, resulting in secondary injury mechanisms that lead to irreversible neuronal death and, ultimately, various degrees of neurological impairment and paralysis (Ahuja et al., 2017). Secondary injury encompasses multiple processes, including disruption of the blood-brain barrier, local ischemia, axonal shearing, gliosis, and neuroinflammation (Anjum et al., 2020; Sterner and Sterner, 2022). In recent years, preclinical research advances have significantly contributed to elucidating the pathology driving secondary injury mechanisms and identifying potential therapeutic targets. However, these findings have consistently failed to translate into effective therapies that can restore function in people living with SCI. The reason for this failure are multiple and are often common to other neurological/neurodegenerative disorders. Rodent models, which are widely utilized in SCI research, lack consistency and reproducibility in the way injury is induced within and across laboratories, and in how functional outcomes are measured, weakening the foundation upon which translational and clinical studies are designed. This may lead to negative or inconclusive data that cause potentially effective drugs to be dismissed or to an overestimation of efficacy that pushes unsuitable drugs into clinical trials. While it is impossible to entirely mitigate the variability of injury induction, which is largely dependent on the skills of the operator, the assessment of functional outcomes can be improved by developing unbiased, investigator-independent approaches.

Since 1995, the Basso Beattie Bresnahan (BBB) scale (Basso et al., 1995) and later the Basso Mouse Scale (BMS) have been the gold standard for assessment of locomotor function after SCI in rats and mice, respectively (Basso, 2004). Based on the visual evaluation of rodent behavior in the open field by two examiners positioned opposite to each other, BBB and BMS are easy to execute and reliable methods to assess locomotion that can be repeated with minimal stress to the animals, thus being valuable for tracking locomotor recovery over time. However, as with other behavioral tests, these tests have major limitations in that they are subjective and have limited sensitivity in detecting subtle changes in specific aspects of movement that go unnoticed by the human eye. As a result, unless changes in locomotor performance are substantial, BBB and BMS may not be able to identify them.

To improve investigator accuracy and bias in behavioral tests, video recordings have been implemented (Gomez et al., 2018; Sparrow et al., 2017). Nevertheless, extracting discrete behavioral metrics using observers is labor-intensive, time-consuming, and error-prone. Instead, supervised learning methods in which an “action classifier”, or machine learning algorithm, is trained to recognize specific patterns defined by researchers is a much more effective approach to detect changes in behavior while minimizing user-dependent variability and bias, and increasing sensitivity, accuracy, and data output (Anderson and Perona, 2014). The classifier undergoes iterative training and evaluation until it attains reliable metric (e.g. stride length and limb angle) estimations. Within this concept, we developed two separate marker-less kinematic analysis paradigms in mice, MotorBox (MB) and MotoRater (MR), based on deep-learning algorithms generated with the DeepLabCut (DLC) open-source toolbox. DLC employs object recognition and semantic segmentation algorithms derived from pre-trained ResNets and deconvolutional layers to achieve robust pose predictions that surpass commercial solutions (Lauer et al., 2022; Mathis et al., 2018), (Sturman et al., 2020). Current implementations of DLC have facilitated a significantly broader exploration of hindlimb locomotor deficits in conditions such as SCI and experimental autoimmune encephalomyelitis (EAE), a model of multiple sclerosis (Aljovic et al., 2022; Eisdorfer et al., 2022; Sato et al., 2022). However, these approaches are limited to a few parameters, and focus only on the analysis of a single hind limb. Here, we validated MB and MR by comparing them with the traditional BMS evaluation and extracted metrics of movement and gait that can provide an accurate and sensitive representation of mouse locomotor function post-injury, while eliminating investigator bias and variability.

## 2. Materials and Methods

### 2.1. Mice

Wild type C57Bl/6J female mice aged 8 weeks were used for these studies. Mice were group-housed (maximum 5 mice/cage) in the Animal Facility of The Miami Project to Cure Paralysis, University of Miami Miller School of Medicine, in a virus and antigen-free, temperature-controlled environment with a 12 h light/dark cycle. Mice had access to food and water ad libitum. After surgery, mice were individually housed and tested longitudinally (from the day before injury to 35 days post injury) with BMS, MB and MR on the same day. All experiments were performed in accordance with the guidelines of the Institutional Animal Care and Use Committee of the University of Miami Miller School of Medicine under approved protocols.

### 2.2. Spinal Cord Injury (SCI)

Moderate contusive spinal cord injury (SCI) was performed with the Infinite Horizon (IH) device (model #0400) according to established protocols (Brambilla et al., 2005; Tsenkina et al., 2015). Briefly, after anesthesia with isoflurane (induction = 3%; maintenance = 2%, combined with 1.5 l/min oxygen), mice were shaved over the thoracic spine, placed in a stereotaxic frame, and laminectomized between vertebrae T7 and T9. Contusion injury was induced by lowering the vertical shaft of the IH impactor over vertebra T8 at a force of 50 kilodynes. Impact force was measured with a pressure transducer. After surgery, mice were single housed and administered subcutaneous Ringer’s solution to prevent fluid loss, gentamicin (2-5 mg/kg) to prevent urinary tract infections, and buprenorphine (0.5-1.0 mg/kg) for pain relief. Manual bladder expression was performed twice a day until mice spontaneously regained function.

### 2.3. Basso Mouse Scale (BMS)

The Basso Mouse Scale for locomotion (BMS) is a widely used scale for evaluation of locomotion in the open field, specifically designed to assess hindlimb functional deficits in mice (Basso et al., 2006). In this test, each mouse is individually placed in a square box measuring 2 ft x 2 ft and allowed to freely move for a period of 4 minutes. Movement of each hindlimb is assigned a 1 to 9 score based on its degree of locomotor function, where 0 = complete paralysis, 1 = slight ankle movements on either side, 2 = extensive ankle movement on either side, 3 = plantar placement with or without weight support, or occasional dorsal stepping without plantar stepping, 4 = occasional plantar stepping without forelimb to hindlimb coordination, 5 = frequent plantar stepping with no coordination, or with some coordination along with rotated paws at initial contact and lift off, 6 = frequent plantar stepping sometimes or mostly coordinated, with either parallel paws at initial contact or rotated paws at initial contact and lift off, 7 = frequent plantar stepping that is mostly coordinated, with parallel paws at initial contact and lift off, and possibly severe trunk instability, 8 = same descriptors as score of 7 but with mild trunk instability or normal trunk stability, and with either tail down or going up and down, 9 = normal hindlimb locomotion with consistent coordinated plantar stepping, with parallel paws, normal trunk stability, and tail always elevated. Mice were assessed with the BMS prior to SCI to establish a baseline level of locomotor function, then at 1, 7, 14, 21, and 28 days post injury (dpi) to monitor hindlimb functional performance overtime.

### 2.4. Kinematic analysis with the MotorBox

The MotorBox (MB) is a system designed to assess mouse locomotor function and kinematics in the open field in an unbiased manner taking advantage of the deep-learning algorithm DeepLabCut (DLC), an open-source toolbox for markerless pose estimation (Mathis et al., 2018; Nath et al., 2019).

#### 2.4.a MotorBox apparatus

The custom-made MB apparatus consists of a transparent square box (length = 35 cm; width = 35 cm; height = 40 cm) constructed with four acrylic panels (panel thickness = 0.5 cm) attached to an acrylic floor base (length = 43 cm; width = 43 cm; thickness = 1 cm) (Fig. 1A, B). The chamber is elevated above the ground thanks to four leg pegs (length = 45 cm; diameter = 2.5 cm) attached, at the top, at each corner of the chamber base and, at the bottom, at each corner of the floor platform. At the center of the floor platform, a GoPro HERO8 Black camera is positioned and held in place in a bottom-up recording mode by a 3D-printed ski boot GoPro mount (Fig. 1C). To minimize film grain and motion blur, video recording parameters are set as follows: 1) Resolution = 4k; 2) Frames per second (FPS) = 60; 3) Lens = Linear; 4) HyperSmooth function = On; 5) Bit rate = High; 6) Protune function = On; 7) Shutter = 1/3840; 8) White Balance = Auto; 9) ISO Min = 100; 10) ISO Max = 6400; 11) Sharpness = High; 12) Color = Flat. To ensure optimal tracking, lighting conditions are set for low shot recordings, where four bi-color 5600k light sources (PLV-S116, Raleno) are positioned at the four corners of the MB chamber base, pointing towards the center to avoid shadows (Fig. 1F). For the purpose of our analysis, mice were video recorded in the MB chamber for 2 minutes and the resulting MP4 video files (approximate size = 800,000 kb) were saved on University of Miami approved cloud storage systems until further processing.

**Figure 1.**
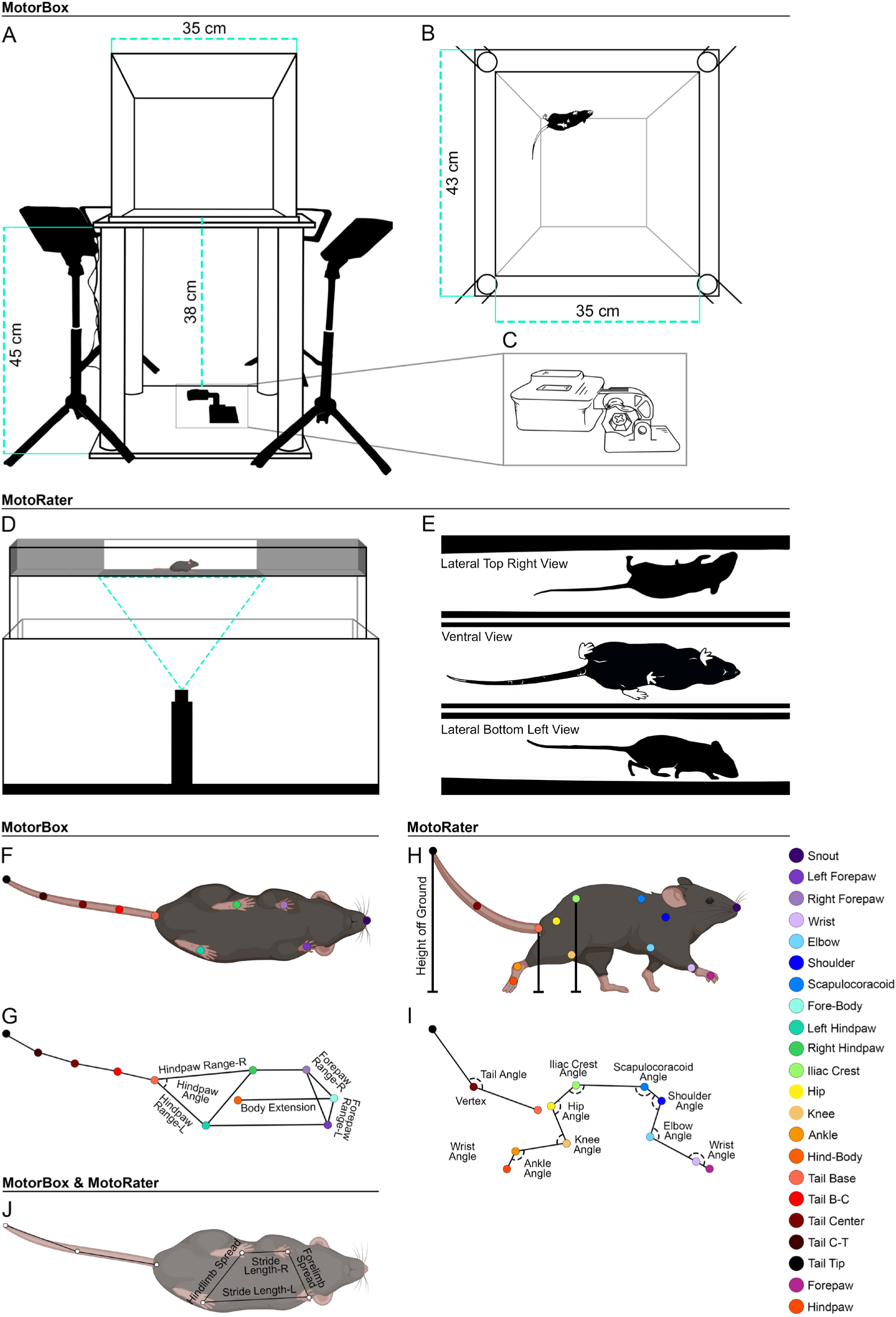
MB and MR equipment and virtual tracking. (A) Schematic of the MB chamber, with camera placement and lighting setup. (B) Field of view from the bottom where DLC tracking occurs within the MB apparatus. (C) 3D printed ski boot mount for positioning of GoPro camera. (D) Schematic of the MR linear runway, with camera placement and recording filed. (E) The lateral and ventral views form the MR. (F) Virtual label placement for DLC tracking in the MB. (G) Skeleton schematic representing unique metrics assessed with the MB. (H) Virtual label placement for DLC tracking in the MR. (G) Skeleton schematic representing unique metrics assessed with the MR. (J) Metrics and virtual marks overlapping in MB and MR.

#### 2.4.b Testing protocol

To fully capture the exploratory behavior of the animals in the open field, mice were not subjected to a habituation period in the chamber prior to testing. On test day, all lighting sources were turned on prior to transferring the mice to the testing room. Each mouse was placed at the center of the MB chamber and video recorded for 2 minutes while moving freely. Throughout this time, the experimenter stepped away from the apparatus to avoid interference with mouse performance. Prior to testing the next animal, the apparatus was thoroughly cleaned with water.

#### 2.4.c Automated kinematic analysis with DeepLabCut (DLC)

All analyses were performed using the Anaconda environment provided by the Mathis lab (Mathis et al., 2018). For this project, a workstation was configured with a PNY GEFORCE RTX 4080 GPU, an Intel i9 9900K CPU, and 64 GB of RAM using Windows 10. The chosen versions for the TensorFlow framework and GPU acceleration include CUDA 12.0, NVIDIA driver 527.37, Python 3.8.12, TensorFlow 2.10.1, and DLC 2.3.0.

Automated kinematic analysis was performed by processing the obtained videos through DLC. Prior to applying DLC, MP4 files underwent cropping to encompass solely the regions where the mouse was in active exploration. To enable pose estimation, specific body parts, namely nose, each of the four limbs, top and bottom of the body, and tail, were marked with “virtual labels” (Fig. 1F, G, J). The tail was labeled with five equidistant marks, from the base to the tip (Fig. F, G, J). Marked body parts allowed to gather kinematics data descriptive of the mouse range of motion, speed, and agility. Prior to its implementation in the kinematic analysis of SCI mice, the DLC algorithm was trained with videos from mice exhibiting various degrees of locomotor deficits and paralysis in different lighting conditions to minimize overfitting. From the training videos, selected frames were extracted using the automatic extraction method, then processed with the *kmeans* clustering algorithm using a cluster step of 1 and cropping frames disabled. This algorithm works by dividing a set of *n* observations in *d*- dimensional space (**R***^d^)* into *k* clusters to minimize the *squared-error distortion* or the mean squared distance between each data point and its closest center (Tapas Kanungo. David M. Mount, 2002). Three hundred and seventy frames were labeled from the selected training videos using DLC’s Napari GUI plugin. From those, a training dataset was created using five different network architectures: ResNet-50, ResNet-101, ResNet-152, MobileNet-v2-1.0 and EfficientNet-b6. ResNet-152 was determined to provide the best training outcome. This network is part the Residual Networks (ResNet) repository, consisting of networks capable of integrating features of varying importance and classifiers in a multilayer fashion that can be enriched through an increase in stacked layers. ResNet-152 is the network architecture with the largest depth of layers for image recognition (Sun, 2016). The training dataset was then augmented using Imgaug (Mathis et al., 2020), a library for image augmentation, that creates a modified version of the dataset as a means of introducing diversity. This method helps minimize overfitting and improve model accuracy. After defining the training parameters, the network underwent training for 200,000 iterations. The training process continued until the losses reached a plateau at 0.001 by 25,000 iterations. Following the training phase, the network was evaluated until the pixel error reached satisfactory levels of 3.25 for the training error, and 6.93 for the testing error. Predictions above 0.6 were filtered, and a short window (n=5) interpolation was performed on all values. Ultimately, the algorithm was fully trained, and batch processing was implemented for all the videos. Our DLC workflow is publicly available in the GitHub repository (https://github.com/MaureenAscona/DLC-MotorBox/tree/main)

#### 2.4.d Outcome measures

Once the cropped video files were uploaded into DLC, they were processed to obtain x and y coordinates of all virtually labeled body parts as the animal moved. These data, output in .csv file format, were then processed through Python packages to extract a variety of outcome measures representative of kinematics patterns and movement characteristics. In the equations used to calculate the various outcome measures, the coordinates of the various body parts were labeled as follows: left forepaw (x1, y1), right forepaw (x2, y2), top body (x3, y3), left hindpaw (x4, y4), right hindpaw (x5, y5), bottom body (x6, y6), tail base (x7, y7), tail b-c (x8, y8), tail center (x9, y9), tail c-t (x10, y10), tail tip (x11, y11).

##### Limb Spread

“Limb spread” refers to the distance between forelimbs or hindlimbs throughout the exploration period. Spread is calculated as the distance, at each frame, between the position of front left and right paws, or back left and right paws. It represents the distance between the two limbs and was calculated using the formula:

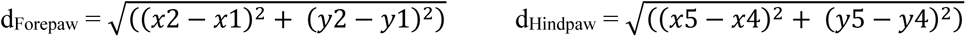

##### Stride Length

“Stride length” refers to the distance between fore- and hindlimbs on the same side of the body. Stride length is calculated as the distance, at each frame, between the position of front and back right paws, or front and back left paws. The stride length was calculated using the formula:

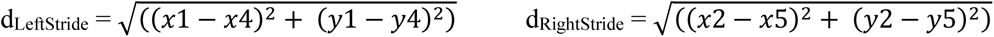

##### Synchronicity

“Synchronicity” describes a quadrupedal gait where the inter-paired appendages exhibit synchronous phasing, meaning that the timing of movements between forelimbs and hindlimbs is coordinated (Struble and Gibb, 2022). In this phased gait, the limbs within the same girdle (forelimbs or hindlimbs) are out of phase with each other, ensuring that there is always at least one limb serving as a support during locomotion. Synchronicity can be extrapolated from stride length and spread measures. In a synchronous gait, short and long stride lengths alternate in a consistent pattern, paired with a consistent distance between forelimbs and hindlimbs (limb spread). These measurements serve as indicators of synchronous movement during locomotion.

##### Paw Range

“Paw range” identifies the position and extension of a limb in relation to a reference point. It serves as a surrogate measure to determine the swinging motion and weight-bearing of the limb during movement. Hindpaw range is determined by measuring the distance between a hindpaw and the base of the tail (Fig. 1G). To calculate this distance over time, the distance formula is employed using the coordinates of the left hindpaw (x4, y4) and of the tail base (x7, y7). Forepaw drag is assessed by measuring the distance between a forepaw and the “top body” (x3, y3), which is identified as the base of the neck (Fig. 1G).

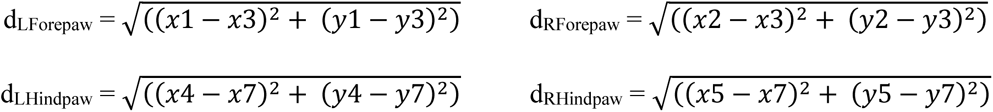

##### Body extension

“Body extension” refers to the distance between the top (x3, y3) and the bottom part (x6, y6) of the body. The top is identified as the base of the neck, and the bottom refers to the lower abdomen, at the insertion point of the hindlimbs (Fig. 1G). This metric provides insights into the degree of hunching or extension exhibited by the mouse at any given moment. This parameter is calculated by applying the distance formula between top and bottom body at each frame.

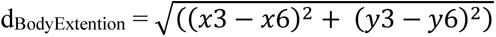

##### Speed, Velocity, and Acceleration

These are key parameters to evaluate mouse movement and can be calculated for each individual body label. To measure overall movement, the tail base label is used, as this mark has the highest probability to be in frame and is the least prone to mislabeling. To determine the movement of each limb, the left and right fore- and hindpaw labels are used, and for tail speed, the tail tip mark is used.

*Speed* is calculated based on distance and time, where distance is obtained by the position of the individual body label in the current frame compared to the previous frame, and time is determined from the current frame divided by the number of frames per second. A conditional restriction was implemented to exclude instances where the speed of the tail base was below 0.5 cm/s, serving as a filter to eliminate periods when the mouse was stationary.

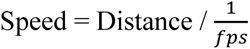

*Velocity* assesses the speed of the mouse in a particular direction and is calculated as the change in only the x coordinate of a body label, or it’s displacement, across successive frames over time.

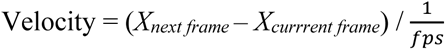

*Acceleration* measures how rapidly mice altered their velocity according to the formula:

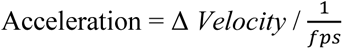

##### Hindpaw Angle

“Hindpaw angle” is calculated by using three distinct points: left and right hindlimbs and tail base. The hindpaw angle measures the clockwise angle between 2 vectors, the tail base to the left hindpaw and the tail base to the right hindpaw, respectively (Fig. 1G, Hindpaw Angle). This angle provides insight into the hindlimb range of motion of the animal at each frame. The formula used to calculate the angle is:

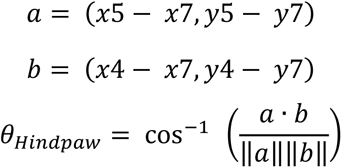

##### Tail Position

This metric assesses the tail’s height off the ground based on the length of the tail. Since the animal is observed from a bottom-up perspective, the tail height off the ground is estimated based on the length of the tail, wherein a longer tail length is reflective of a resting or flaccid tail, while a shorter tail length refers to an upright tail. To calculate the tail length, several distances are determined using the distance formula, which include the distance between the tail base (x7, y7) and tail b-c, tail b-c (x8, y8) and tail center, tail center (x9, y9) and tail c-t, and tail c-t (x10, y10) and tail tip (x11, y11) (Fig. 1G). These distances are calculated at each frame, and their sum provides the total tail length.

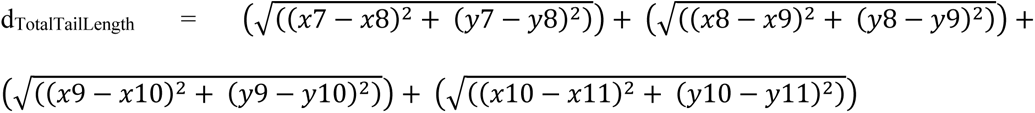

##### Open field exploration

In addition to providing accurate kinematics analytics, the MB also serves as a platform to assess open field exploration in mice. To achieve an accurate and unbiased analysis of exploration times and patterns, the MB chamber is subdivided into five regions, with a center zone (21 cm x 21 cm) surrounded by four corner zones. As with a traditional open field apparatus, the time spent in each region of the chamber is measured to gain insights into the overall level of locomotor activity as well as anxiety-like behavior. The movement patterns of each body part and the mouse walking trajectory can be tracked and graphed by plotting the x,y coordinates of the specific body part in each video frame. The x and y coordinates are established using the DLC infrastructure, where the origin (0, 0) is positioned at the top-left corner of the video. Coordinate values increase moving towards the bottom-left and top-right corners of the frame (Fig. 1B).

### 2.5. Kinematics analysis with the MotoRater

#### 2.5.a MotoRater apparatus

The MotoRater (MR) is a device manufactured by TSE Systems that assesses kinematics in rodents. For our study, we used the configuration for mice, which consists of a narrow plexiglass runway (length =150 cm length, width =13 cm, height = 40 cm) that forces the animal to walk in one direction. As the animal walks, its movements are recorded with a high-frame rate camera positioned underneath the runway and pointing upward (Fig. 1D). The side views of the mouse were captured thanks to two mirrors placed at an angle along the length of each side of the runway (Fig. 1E). To optimize lighting conditions for our recordings, we modified the apparatus by placing an additional light strip at each side of the runway. Although the manufacturer developed the MR apparatus to be used with physical markers to label the animal’s body parts, we applied a DLC based markerless approach for unbiased pose estimation, similar to what we implemented for the MB apparatus (Fig. 1D, E, H).

#### 2.5.b Testing protocol

Before SCI, mice underwent a habituation period in the MR apparatus to familiarize with the environment and minimize anxiety-related behaviors that might affect task performance on test day. During habituation, mice were positioned at the center of the runway and allowed to walk towards a black box placed at the end of the runway and containing litter from their home cage. This served as incentive for the mice to walk across the runway. If the mouse successfully returned to the black box, it was placed back on the runway at a gradually increased distance from the box. This process was repeated until the mouse could walk uninterrupted from one end of the runway to the opposite end where the black box was positioned. On test day, the mouse was recorded as it walked across the entire runway. Of note, mouse gait cycles were analyzed only in the “locomotor football pattern”, defined as symmetrical intra-paired appendage phased gating (Hildebrand, 1977; Struble and Gibb, 2022).

#### 2.5.c Automated kinematic analysis with DeepLabCut

TSE Motion software was used to collect videos at a frame rate of 200 fps and a resolution of 1600 x 1400 pixels. Instead of the TSE Motion software provided by the manufacturer, we analyzed the videos with a DLC-based approach, similar to that implemented for the MB, which allowed to generate over 800 distinct kinematics measurements. TSE Motion software outputs .avi files whose starting and ending positions can be trimmed. We trimmed the videos to select only the footage where the entire animal, from tail tip to nose, was within the frame at each observation (ventral, lateral right/left) (Fig. 1E). For the ventral view, the Unique body parts annotation was selected as the ventral view contains fewer individual landmarks required for tracking (Fig. 1H). As described in section “Automated kinematic analysis with DeepLabCut (DLC)”, and using the same environment as the MotorBox, several different videos were selected for algorithm training: under different lighting conditions (with/without light strips), with mice at different stages of disease, and different behaviors. For this training, 214 frames were extracted from 25 different videos using the automatic extraction method and labeled using Napari. A multi-animal training dataset was created using resnet-152, and the network was trained for 20,000 iterations, when losses plateaued at 0.0004 with an acceptable training error of 3.19 px and testing error of 4.69 px. The DLC workflow for the MR apparatus can be found in this GitHub repository (https://github.com/MaureenAscona/DLC-MotoRater).

#### 2.5.d Outcome measures

Some of the metrics obtained with the MR overlap with those obtained with the MB described above. As the MR is based on a bottom-view recording, stride length and limb spread were analyzed as described for the MB above. Additionally, since the MR allows for a side view of the animal during movement, this uniquely allowed for analysis of fore- and hindlimb range of motion based on joint and limb angles. Collected videos were imported into DLC, then csv files containing pose predictions of all specified body points were exported and converted into pandas data frames. Various Python packages were used to calculate the desired outcome measures at each frame. This workflow can be found in this GitHub repository (https://github.com/MaureenAscona/DLC-MotoRater).

In the equations used to calculate the various outcome measures, the coordinates of the various body parts were labeled as follows: top right/bottom left forepaw (x1, y1), top right/bottom left wrist (x2, y2), top right/bottom left elbow (x3, y3), top right/bottom left shoulder (x4, y4), top right/bottom left scapulocoracoid (x5, y5), top right/bottom left iliac crest (x6, y6), top right/bottom left hip (x7, y7), top right/bottom left knee (x8, y8), top right/bottom left ankle (x9, y9), top right/bottom left hindpaw (x10, y10), top right/bottom left tail base (x11, y11), top right/bottom left tail center (x12, y12), top right/bottom left tail tip (x13, y13), center left forepaw (x14, y14), center right forepaw (x15, y15), center left hindpaw (x16, y16), center right hindpaw (x17, y17), center tail base (x18, y18), center tail center (x19, y19), center tail tip (x20, y20).

##### Oscillation Pattern

To assess the “oscillation pattern”, mice are made to walk on the runway in one uninterrupted pass, meaning that if the animal mouse hesitates, or pauses to sniff, scratch, or groom the walk is restarted as the pass is considered incomplete. This stringent exclusion criterion is set to allow for the most accurate representation of the walking oscillation pattern of the mouse. This oscillation pattern is obtained by graphing the x coordinates of the 4 limbs over the span of the recording, which in a healthy mouse shows synchronous phasing of inter- paired appendages (Struble and Gibb, 2022).

##### Forelimb Angles

Forelimb angles were tracked across the length of the video using 3 coordinate points. Wrist, elbow, shoulder, and scapulocoracoid angles of left and right sides were measured (Fig. 1I). Angles were calculated as described above in “*Hindpaw Angle*” by calculating the vector from the point of origin (wrist), in this case the wrist angle, to point A (hindpaw), and the vector from the point of origin to point B (elbow). The “atan2” function was then used to calculate the angle vertex in radians. The angle was then converted to degrees and the internal angle was calculated.

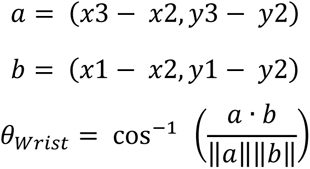

##### Hindlimb Angles

Hindlimb angles were assessed at each frame using three coordinate body points, with the vertex containing the desired angle. The Iliac crest, hip, knee, and ankle angles of left and right sides were measured (Fig. 2D). Angles were calculated as described above in “*Forelimb Angles*”.

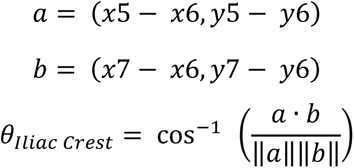

**Figure 2.**
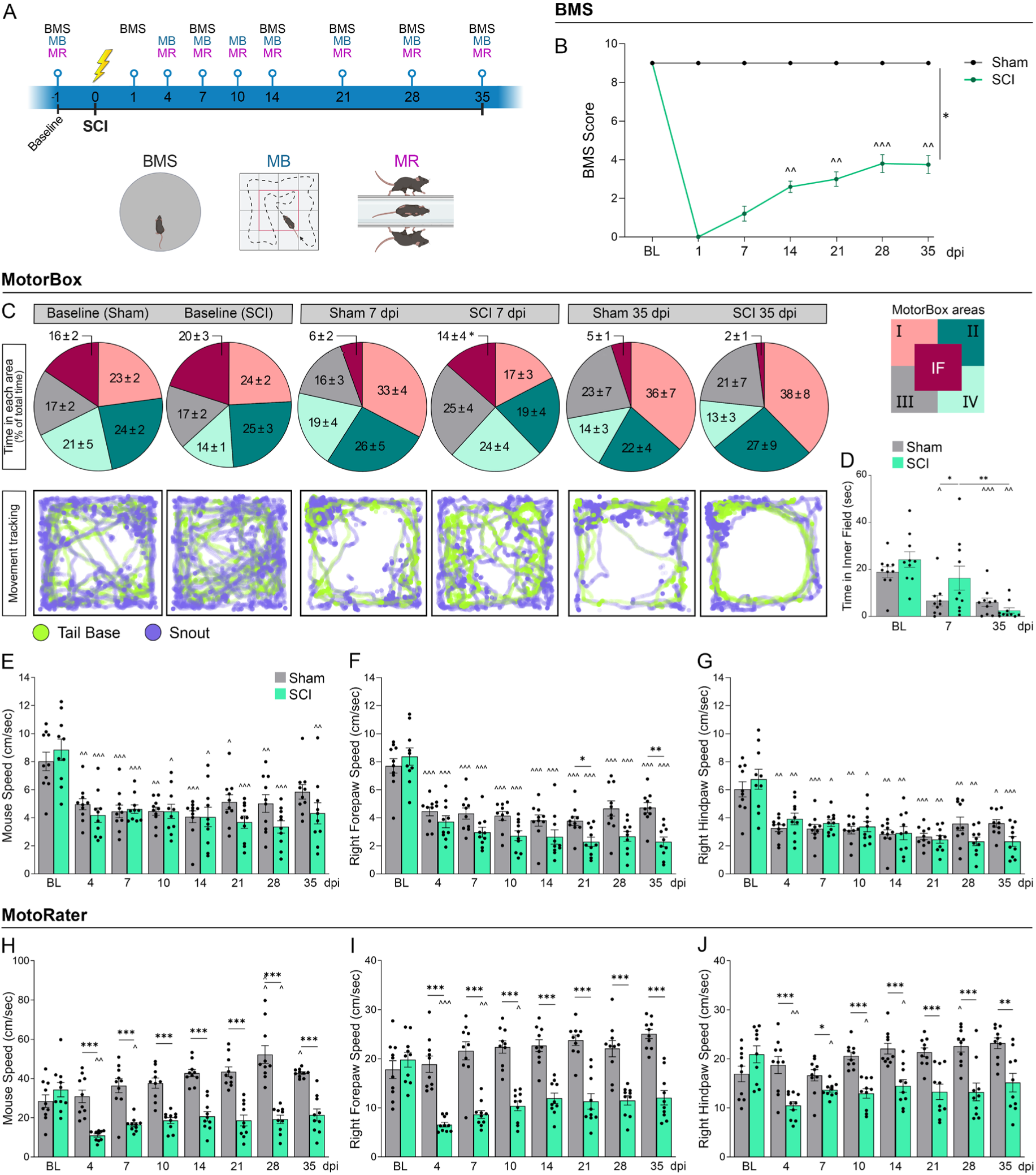
Detection of SCI-induced changes in exploration patterns and ambulation metrics. (A) Experimental timeline for behavioral testing of sham and SCI mice. (B) Evaluation of locomotor function overtime with BMS. (C) Analysis of mouse exploratory behavior with the MB at baseline, 7 and 35 dpi in sham and SCI mice; top panels: time spent in each area of the apparatus; bottom panels: mouse tracking using tail base ad snout as virtual marks. (D) Time spent in the inner field (IF) of the MB at baseline (BL), 7 and 35 dpi. (E-G) Speed metrics over time measured with the MB: (E) mouse speed (tail base used as reference mark), (F) right forepaw speed, and (G) right hindpaw speed. (H-J) Speed metrics overtime measured with the MR: (H) mouse speed (tail base used as reference mark), (I) right forepaw speed, and (J) right hindpaw speed; n=10/group. *p≤0.05, **p≤0.01, ***p≤0.001 between sham and SCI. ^p≤0.05, ^^p≤0.01, ^^^p≤0.001 compared to BL, Repeated Measures Two-Way ANOVA.

##### Tail Angle

The tail angle metric is used to assess tail tone and curvature over time. It is calculated as described above with the tail center as vertex, the tail base as point A, and the tail tip as point B. An angle that approaches 180° indicates a straight up tail from base to tip, whereas an acute tail angle indicates a tail curved towards the center. This metric provides insights into tail flaccidity and muscle impairment in CNS injury.

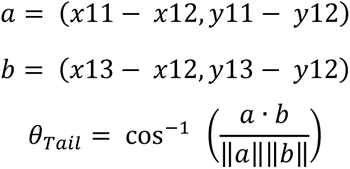

##### Height off Ground

Height off the ground refers to the distance of specific body points from the floor. Tail tip, tail base, iliac crest, hindpaw and forepaw heights are assessed from both the left and right lateral views. This metric is calculated by finding the minimum y value of the hindpaw (*y constant*) throughout the length of the recording, then subtracting it by the y value of all other body points over time (Fig. 1H).

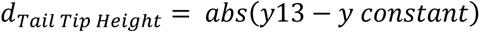

##### Speed, Velocity, Acceleration

Speed, Velocity, and acceleration were calculated as described in the “*Speed, Velocity, and Acceleration*” MotorBox section above, where the speed was calculated as the distance traveled at each frame, the velocity was calculated as the displacement of the x coordinate traveled over time, and the acceleration was calculated as the delta velocity divided by the time between consecutive frames. Mouse speed was calculated using the tail base on the left lateral view of the animal, although, in this case either the left or right lateral view is sufficient. Unilateral fore- and hindpaw speed, velocity, and acceleration were calculated using the respective lateral view to incorporate a 2-dimensional perspective for higher accuracy, as opposed to the MotorBox which is only able to calculate speed, velocity, and acceleration through a 1- dimensional perspective from a ventral view.

### 2.6. Statistical Analysis

Metrics were calculated at each frame and descriptive statistics comprising mean, median, maximum, minimum, and standard deviation were collected from each video. For the MB, outliers were identified through a rolling distance calculator, where a measure was considered an outlier if the distance between the current body point and the point of the next frame exceeded the total length of the mouse. For the MR, outliers were identified using the Hamper filter in combination with any y values whose Z score differed from threshold by more than 3 standard deviations. To determine changes between sham and SCI animals over time, a Two-Way Repeated Measures ANOVA was run and p values equal to or below 0.05 were considered significant. Sample size was established using the G*power 3.1.9.7 calculator, with an effect size set between 1.5 to 2.4 based on preliminary results, alpha value of 0.05, and power of at least 80%. A sample size of 10 animals per group was calculated as more than sufficient for accurate analyses.

## 3. Results

### 3.1. MotorBox allows for sensitive detection of SCI-induced changes in exploration patterns

To explore the potential of MB and MR testing in parsing out injury-induced changes in locomotion, we analyzed mice with the MB and MR in parallel with BMS evaluation, the gold standard for assessment of mouse locomotion after SCI. BMS assessment was performed 1 day prior to surgery (baseline, BL) and at 1, 7, 14, 21, 28, and 35 days post-injury (dpi). Mice were tested with the MB and MR on the same days (except day 1), with the addition of 4 and 10 dpi (Fig. 2A). Consistent with the published literature from our group and others (Brambilla et al., 2005; Gao et al., 2023), BMS testing revealed spontaneous locomotor recovery at 7 dpi in mice with SCI. This plateaued at 28 dpi at a score of approximately 3.5, which was maintained thereafter, indicating regained plantar stepping (Fig. 2B).

MB testing was employed to examine exploration patterns in the open field by measuring the time spent in specific areas of the apparatus (Inner Field = IF; Quadrant I = I; Quadrant II = II; Quadrant III = III; Quadrant IV = IV) (Fig. 2C). This was enabled by tracking the mouse with virtual marks on the tail base and snout. Exploration patterns provide information on injury-induced anxiety-like behaviors. Compared with their corresponding baseline, sham controls showed a reduction in the time spent in the center of the apparatus (IF) at both 7 dpi (acute time point) and 35 dpi (chronic time point) (Fig. 2C, D). Given that mice have a natural tendency to avoid open areas, this suggests that anxiety/stress levels decrease over time in uninjured animals likely as a result of habituation. In contrast, the time spent in the IF at 7 dpi in SCI mice was similar to that at baseline and significantly higher than in the corresponding sham group (Fig. 2C, D). This is indicative of injury-induced anxiety/stress, which ultimately dissipates by 35 dpi, likely due to habituation. Changes in the exploration patterns were clearly demonstrated by mouse tracking using snout and tail base marks (Fig. 2C). Exploratory patterns could not be tracked with the MR, given that mice only move in one direction in this apparatus.

### 3.2. MotoRater and MotorBox allow for detection of SCI-induced changes in speed and acceleration

The MB and MR tests were conducted in parallel to extract quantitative information on key locomotion parameters, with speed as the primary metric. In the MB, overall mouse speed (tail base used as the reference point) was significantly reduced in both sham and SCI mice compared to the corresponding baseline, with no differences between groups at any time point (Fig. 2E). This evidence suggests that, for this particular outcome measure, repeated testing led to habituation independent of the injury. Notably, changes in the speed of the individual right limbs (right forepaw and right hindpaw marks were used as reference points) were only evident in chronic SCI, with a significant reduction at 21 and 35 dpi in the right forepaw speed of SCI mice compared to sham (Fig. 2F, G). Similar results were observed for the speed of the left limbs (Fig. S1), making this metric moderately useful in discriminating injured mice from control mice at chronic SCI time points. Additional metrics of locomotion, such as acceleration, proved to be highly variable and, thus, not useful for discriminating injury-induced changes (Fig. S1, S2).

Next, we extracted the same locomotion metrics from the MR analysis. In all speed metrics, sham mice did not show any habituation response (speed reduction) over time (Fig. 2H-J). In contrast, they progressively increased their speed from baseline through the 35 dpi time point. This was observed for mouse speed (Fig. 2H) as well as individual limb speed (Fig. 2I, J; Fig. S1), which is opposite to what observed with the MB (Fig. 2E-G). In SCI animals, the speed was reduced compared to baseline at 4 dpi, did not recover over time, and was significantly lower than that in sham mice at all time points (Fig. 2H-J). These data demonstrate that speed metrics from MR are more sensitive in discriminating injury-induced changes than those from MB. Acceleration metrics also followed a similar pattern, whereby sham mice were able to accelerate over time, while SCI mice did not (Fig.S1, S2). This is likely due to multiple factors: first, mice are video recorded from three different angles in the MR (bottom, and two side views); second, exploration and habituation are not a factor in this system due to the unidirectionality of the motion. Notably, hindlimb speed was similar to forelimb speed in SCI mice in both MR and MB, despite thoracic injury affecting hindlimb function. A possible explanation is that mice adjust the speed of the forelimbs to match that of the impaired hindlimbs to permit coordinated or semi-coordinated movement.

### 3.3. MotorBox and MotoRater allow for sensitive detection of SCI-induced changes in range of motion and joint/limb angles

We next examined the range of movement by observing the net limb motions during the lift and stance phases (Catavitello et al., 2018). In the MB, the range of motion of both hindlimbs (hindpaw marks used as reference points), defined as the distance between the tail base and left or right hindpaw (Fig. 3A, B), remained constant in sham animals, but significantly decreased over time in SCI animals starting at 7 dpi (Fig. 3A, B). Notably, this was not observed in the forelimbs, where the range was similar between sham and injured mice at all times (Fig. S3A). Additionally, after an acute reduction at 7 dpi, the body extension increased over time after SCI (Fig. S3B). This may reflect a more hunched trunk immediately after injury, followed by increased torso extension as the mice recover. Interestingly, by measuring the hindlimb angle, the MB uncovered transient injury-dependent characteristics, where the hindlimbs of SCI mice had a significantly wider spread (angles greater than 120°) than those of sham mice at 4 dpi (Fig. 3 C, D), but returned to values close to baseline by 7 dpi (not shown). This is indicative of hindlimb spastic/rigid paralysis, which progressively and spontaneously improves over time. The hindlimb range of motion measured with the MR produced results highly consistent with those of the MB, making both MB and MR well-suited systems to identify SCI-induced changes in these metrics (Fig. 3E, F).

**Figure 3.**
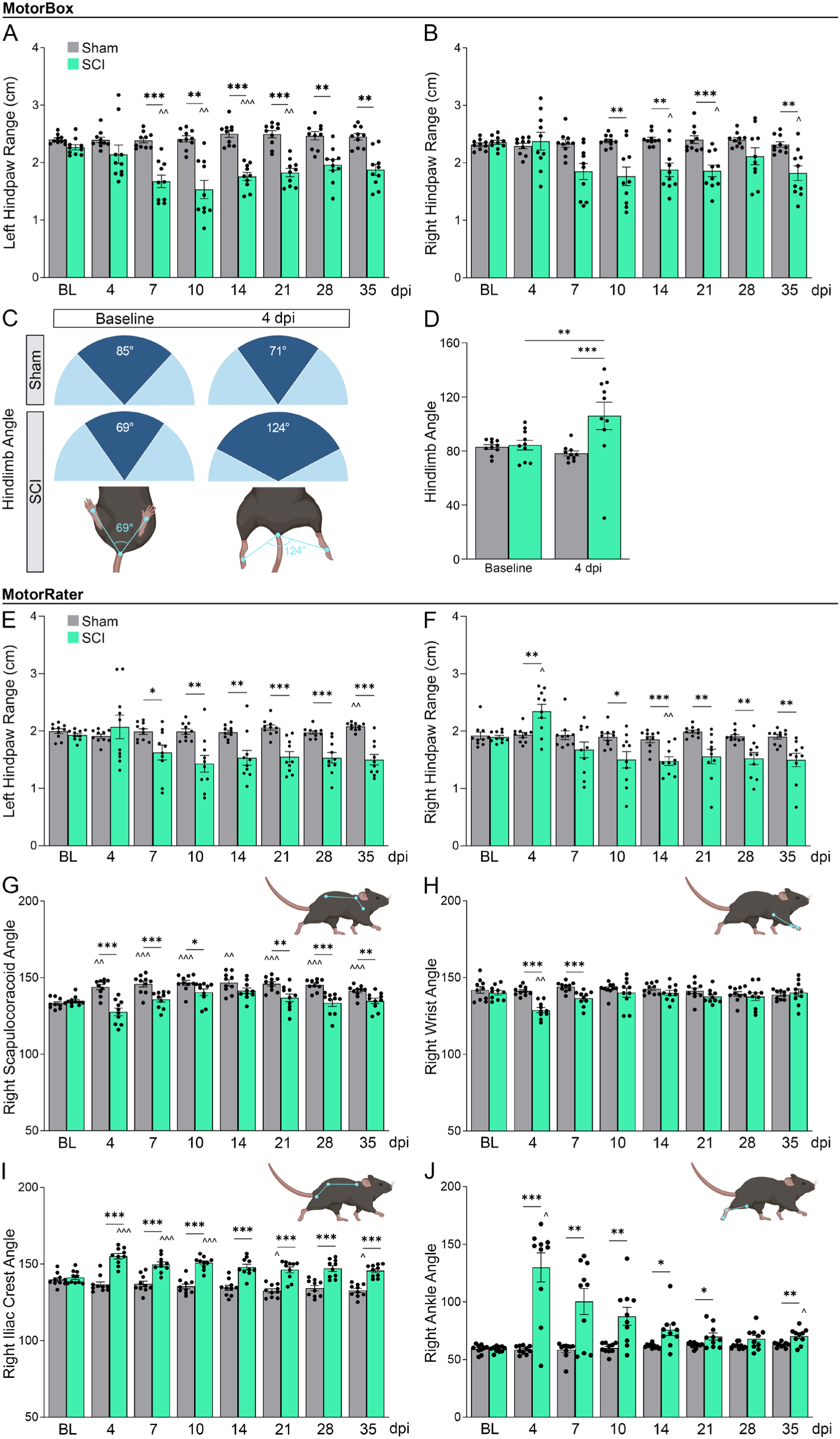
Detection of SCI-induced changes in range of motion. (A, B) Range of motion measured as distance from left hindpaw to tail base (A), and right hindpaw to tail base (B) with the MB. (C) Diagram of the hindlimb angle measured from the two hindpaws to the tail base at BL and 4 dpi with the MB; representative animals are shown. (E, F) Range of motion measured as distance from left hindpaw to tail base (E), and right hindpaw to tail base (F) with the MR. (G-J) Joint angles measured over time with the MR: (G) right scapulocoracoid, (H) right iliac crest, (I) right wrist, and (J) right ankle angles; n=10/group. *p≤0.05, **p≤0.01, ***p≤0.001 between sham and SCI. ^p≤0.05, ^^p≤0.01, ^^^p≤0.001 compared to BL, Repeated Measures Two-Way ANOVA.

A unique feature of the MR is that joint angles (shoulder, scapulocoracoid, iliac crest, pelvic joint, knee hinge joint, wrist, and ankle) can be precisely measured using the lateral view afforded by the apparatus. Several of these metrics exhibited SCI-dependent changes (Fig. 3E- 3J; Fig. S3). For example, a consistent reduction in the right scapulocoracoid angle (above injury level metrics) indicated a persistently diminished range on the right-hand side that never recovered (Fig. 3G). This reflects the SCI-induced deficits in pectoral girdle function. The wrist angle, another above injury metrics, showed similar trends, with significant reductions at 4 and 7 dpi (Fig. 3H). Below injury level, joint angles showed opposite changes, with SCI animals showing increases in both the iliac crest (hip joint) and ankle angles, indicating various degrees of rigid paralysis (Fig. 3I, J).

### 3.4. MotorBox and MotoRater allow for sensitive detection of SCI-induced changes in gait and coordinated walking

Synchronicity between the limbs is essential for effective locomotion (Marder and Bucher, 2001). By tracking each limb independently, both the MB and MR can provide information on stride length and limb synchronicity within and between girdles, thus informing on the degree of movement efficiency and coordination. By examining this rhythmic walking pattern, known as gait, we can characterize abnormal gait as occurrences where mice exhibit deviations from coordinated movements. With the MB, we observed an increased stride length on both the left (Fig. 4A) and right sides (Fig. S4A) at all time points compared to sham mice. The spread between the hindlimbs decreased in SCI mice compared to that in sham mice (Fig. 4B), reflecting rigid paralysis of the lower limbs. Notably, the forelimbs showed increased spread throughout (Fig. S4B), possibly because of the increased effort exerted by the hindlimbs to generate movement. Assessments of stride length and paw spread were perfectly replicated by MR analysis, demonstrating remarkable consistency between the two systems, which can be used interchangeably for these parameters (Fig. 4C, D; Fig. S4C, D).

**Figure 4.**
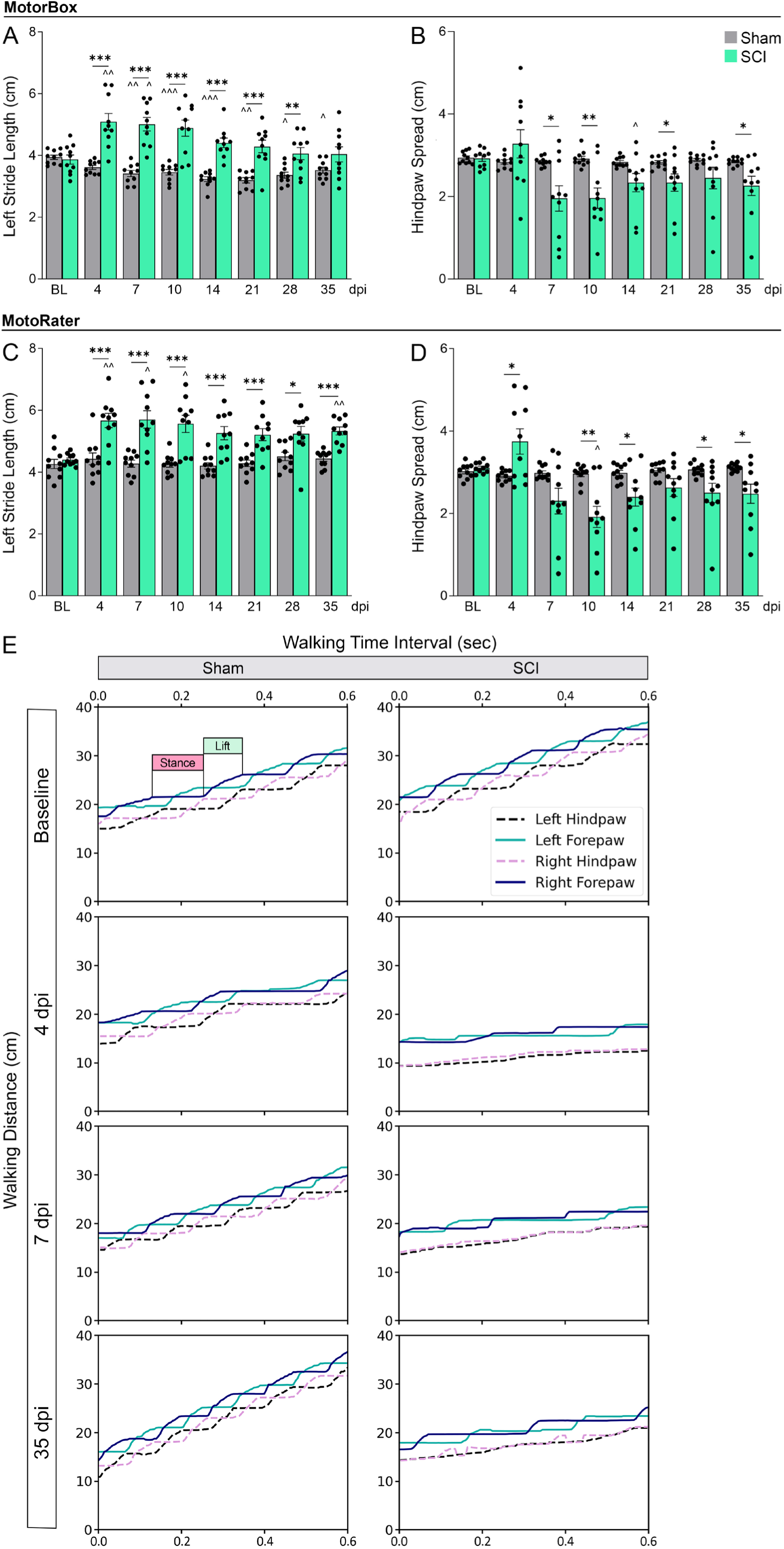
Detection of SCI-induced changes in gait and coordination. (A-C) Movement stride measured as distance from left hindpaw to left forepaw using both MB (A) and MR (C). (B, D) Hindlimb spread measured as distance from left hindpaw to hindpaw using both MB (B), and MR (D). (E) Diagrams illustrating synchronous and asynchronous movement patterns of the four limbs and reciprocal positioning during a 0.6 second travel time in the MR apparatus; n=10/group. *p≤0.05, **p≤0.01, ***p≤0.001 between sham and SCI. ^p≤0.05, ^^p≤0.01, ^^^p≤0.001 compared to BL, Repeated Measures Two-Way ANOVA.

To visualize changes in coordinated walking, with the MR we examined the traces depicting the oscillation patterns of each individual limb during movement (Fig. 4E). Coordination can be visualized by comparing lift and stance, which are defined as the points at which the animal has its paw elevated off the ground and moves across the walkway, and the stage at which the animal has its foot on the ground, respectively. In sham mice, we observed that the left forelimb moved in sequence with the right hindlimb, whereas the right forelimb moved in sequence with the left hindlimb. This is indicative of synchronized stepping or normal gait. In contrast, SCI mice completely lost the ability to perform any stepping in the hindlimbs (below injury level), which caused the forelimbs (above injury level) to elongate their stride and lose synchronized stepping, which was only partially regained at chronic time points (35 dpi) (Fig. 4E).

Synchronous movement was further analyzed with the MR, where different aspects of the lift and stance phases were explored (Fig. 5). Using the y-coordinates of each limb, we determined the number of steps taken and the height of each step as the mouse traveled across the platform., As mice regained motor function in their hindlimbs, we observed that the number of steps they took increased (Fig. 5A), whereas the height (amplitude) decreased (Fig. 5B), indicating that, during recovery, shorter and less effective steps are taken by SCI mice to travel the same distance as the controls. This is exemplified by the travel pattern recorded for each mouse, where the lift and stance phases were tracked over time (Fig. 5C), and each peak depicts a lift off the ground. Movement was quantified by measuring the area under the curve (AUC) of the peaks (Fig. 5E). For accurate quantification of the AUC, it was essential to control for the “fisheye” optical artifact, which results from the fact that the MR utilizes a wide-lens, high-frame-rate camera (Fig. 5C). To eliminate fisheye distortion, post-DLC analyses were implemented, wherein the local minimum/maximum y values were employed to construct a piecewise linear curve. Subsequently, the interpolated curve was subtracted from the original y values for correction (Fig. 5C). After correction, AUC was calculated over time (Fig. 5E). AUCs significantly increased at 14 and 21 dpi, indicating that mice maintained their hindlimbs off the ground for longer periods (Fig. 5D, E;), suggesting hesitation or more lagged intermediate movements. Moreover, we evaluated dragging by analyzing the peaks below 0.20 cm (approximately 3/4 of a mouse’s paw). As anticipated, after SCI, mice spent significantly more time dragging their hindpaws, with the highest dragging time at acute SCI (4 dpi) (Fig. 5F). However, over time, SCI mice demonstrated a notable reduction in dragging times compared to 4 dpi, indicating partial recovery that reinstates lifting behaviors.

**Figure 5.**
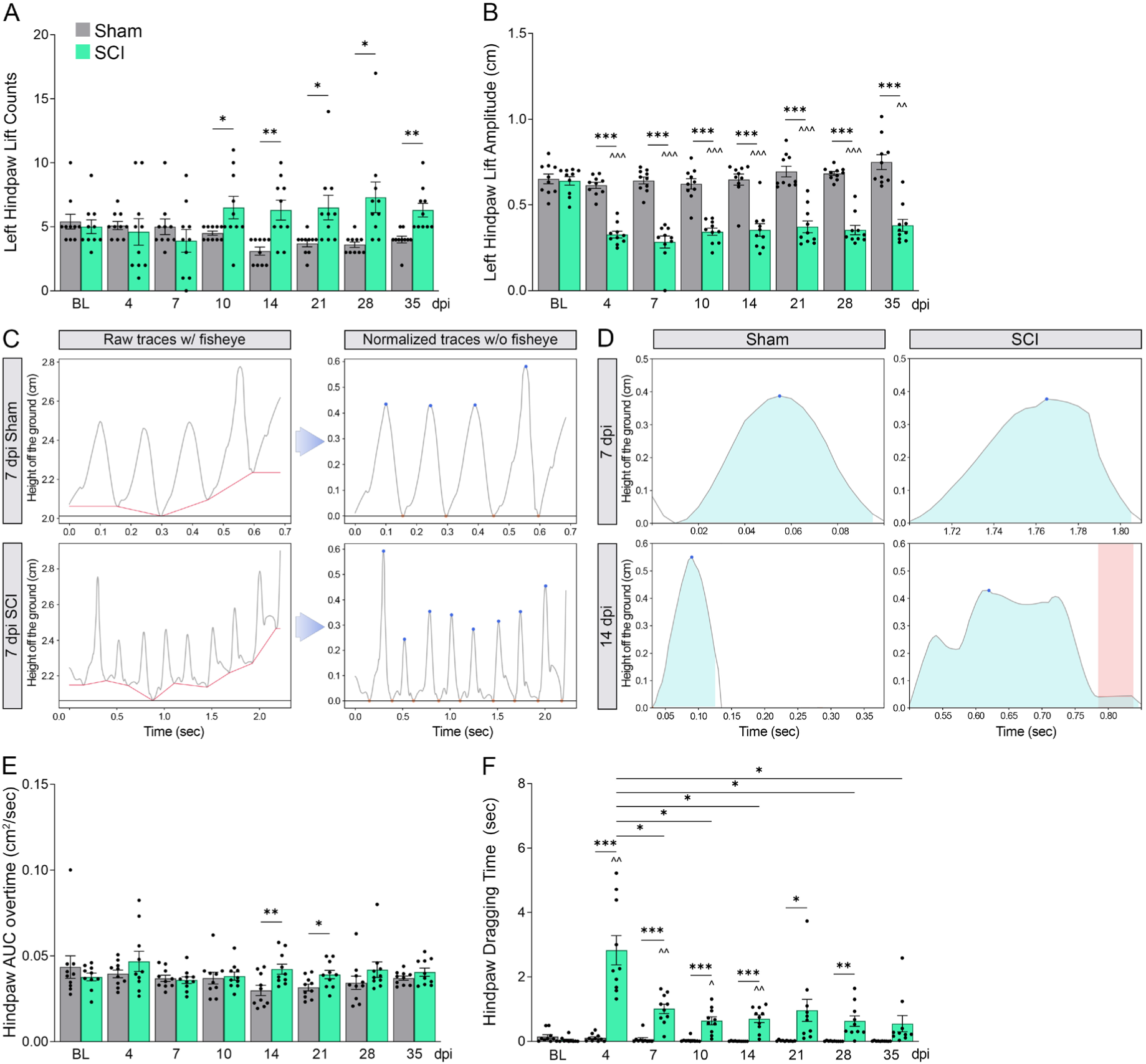
Detection of SCI-induced changes in limb lift dynamics. (A) Lift counts of the left hindpaw overtime assessed with the MR. (B) Lift amplitude (peak amplitude) of the steps taken by the left hindpaw in the MR. (C) Representative diagram illustrating the process of correcting for fisheye distortion and detecting peaks (lift amplitude) and troughs (time in stance) in sham and SC mice at 7 dpi. (D) Representative lift curve shapes at 7 and 14 dpi illustrating the prolonged hindpaw lift off the ground in SCI mice compared to sham. (E) Quantification of the average area under the lift curve (AUC) for both hindpaws in the MR. (F) Time spent dragging the hindpaws in the MR; n=10/group. *p≤0.05, **p≤0.01, ***p≤0.001 between sham and SCI. ^p≤0.05, ^^p≤0.01, ^^^p≤0.001 compared to BL, Repeated Measures Two-Way ANOVA.

### 3.5. Select MB and MR metrics correlate with BMS scores

Given the longstanding use of the BMS as a benchmark for assessing locomotor function after SCI, we performed a correlation analysis between the BMS and each kinematic outcome measure extracted from the MB and MR for SCI mice from 7 to 35 dpi. The correlation strength with the BMS, positive or negative, is depicted as a heat map (Fig. 6). Notably, most MB metrics did not correlate with BMS scores at the acute 7 and 14 dpi time points, when mice were still severely paralyzed (Fig. 6; S5A, B). At later time points (21 to 35 dpi), mostly positive, and a few negative correlations began to emerge. Strong positive correlations were observed, particularly for speed metrics (Fig. 6; Fig. S5E). Acceleration metrics showed a weak to no correlation with the BMS score (Fig. Fig. 6; S5F), and range metrics showed inconsistent correlations.

**Figure 6.**
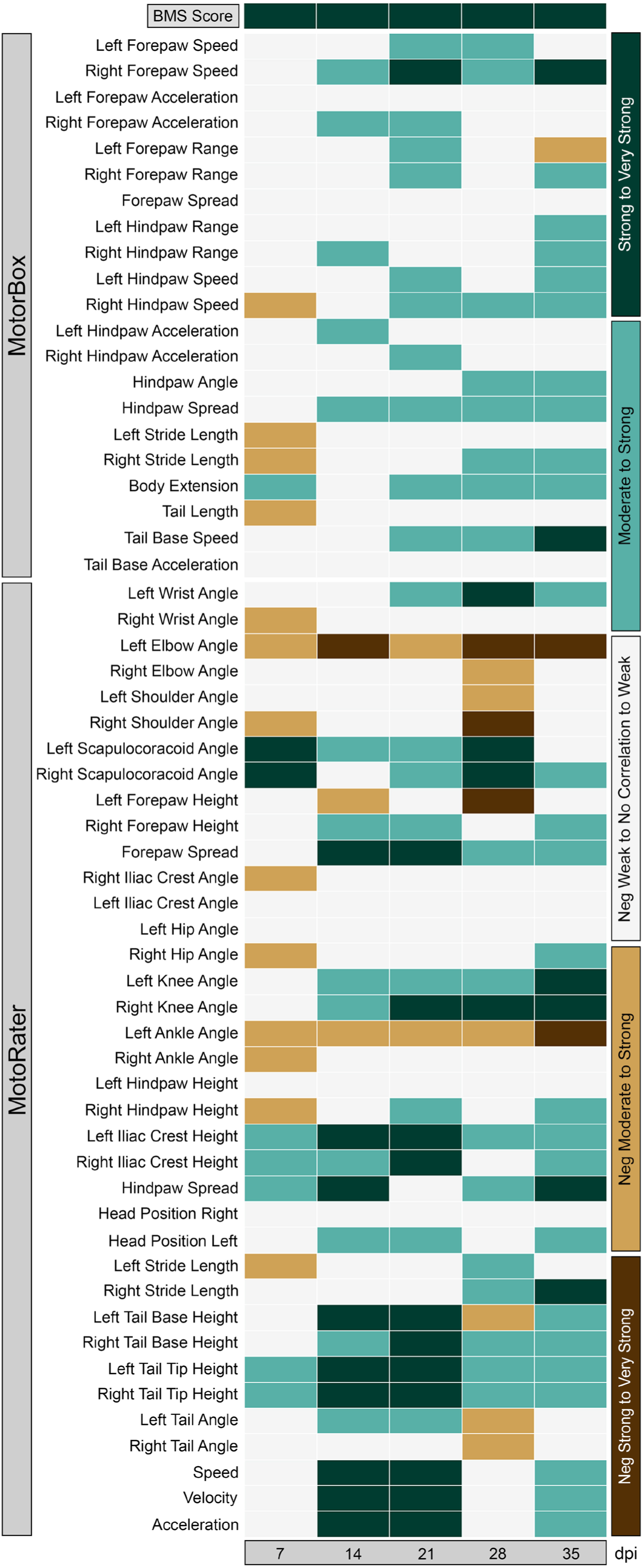
Evaluation Against BMS. (A) Correlation matrix between BMS score and MB and MR metrics calculated with Pearson Correlation at each time point.

Similar to the MB, most MR metrics did not correlate with the BMS at 7 dpi SCI (Fig. 6; Fig. S5D). A few exceptions were found in stance and angle metrics (i.e. iliac height, tail base height, tail tip height, and right scapulocoracoid angle), which were positively correlated with BMS scores (Fig. 6; Fig. S5C). Between 14 and 35 dpi, we observed a generally strong correlation between MR measurements and BMS. Exceptions include some joint angle metrics (e.g. left hip angle, left iliac crest angle, and right ankle angle), which showed no correlation (Fig. Fig. 6; S5H). Wrist and shoulder angles (metrics above injury level) remained negatively correlated throughout recovery. The strongest correlation between MR and BMS was found in speed, velocity, and acceleration metrics, which were not assessed with the BMS. Interestingly, knee angle was highly correlated with BMS scores (Fig. 6; Fig. S5G), however, when analyzing the overall performance of the knee angle metrics in the MR, they seldom demonstrated changes between sham and SCI animals over time (Fig. S3H).

The fact that the BMS correlates with a metric that does not have any utility in depicting SCI-associated deficits further indicates the limitations of the BMS in the accurate, sensitive and robust evaluation of locomotor behavior.

## 4. Discussion and conclusions

We demonstrated that the Motobox (MB) and Motorater (MR) with DCL implementation of pose estimation are accurate, sensitive and unbiased systems for evaluating locomotor function following SCI. We show that the MB is highly accurate in the examination of gait parameters such as stride length and spread, while the MR is especially valuable for detecting differences in ambulation and angular motion, parameters that are not obtained through BMS analysis.

The BMS generates a composite score of multiple observable elements, such as ankle movement, stepping with and without weight-bearing, coordination between girdles, trunk stability, and tail elevation. MB and MR, instead, produce specific quantitative metrics that represent each of these elements, providing precise estimation of stride length, forelimb, and hindlimb spreads as a proxy for coordination, body extension as a proxy for trunk stability, and tail length as a proxy for tail elevation off the ground. Additionally, they include measures of speed, velocity, acceleration, and range of motion (e.g. limb angles) that the BMS cannot provide. Interestingly, our comparative analysis reveals specific advantages of each system.

Constructed from acrylic and featuring relatively compact dimensions, the MB is cost-effective. It can be easily and affordably built with a rapid turnaround time, thus making it a sustainable option for any laboratory. It is also time effective, requiring only one experimenter to place each mouse in the chamber for 2 minutes and upload the videos for analysis, reducing the workload compared with MR and BMS. In addition to kinematics, the MB can also track behaviors indicative of anxiety and stress, which are not obtained with BMS and MR. A caveat of the MB is that, by relying on a solitary ventral view of the mouse, select metrics serve as proxies for certain behavioral characteristics and are not direct measures. One example is tail length, which is used as a substitute for tail height off the ground, thus indirectly estimating tail flaccidity. Moreover, a drawback of the MB is that the absence of a lateral view prevents the exploration of crucial features, such as ankle movements and limb angles, which are partially accounted for in the BMS and directly measured in the MR. Moreover, the 2-minute exploration period may lead to fatigue and disinterest in injured mice, affecting locomotor performance. This may be offset by reducing the exploration time.

MR has distinct advantages over BMS and MB. Its ventral and lateral views provide a clear depiction of the ankle, foot, and tail kinematics, offering a complete portrayal of the mouse’s gait. Similar to the MB, the MR delivers a precise representation of ambulation and synchronization with the added advantage of capturing angular motion and weight-bearing behaviors. These metrics are more sensitive and accurate than the BMS observations. Limb movement synchronization can be quantified by both MB and MR, but the MR, because of the unidirectional linear pattern of motion allows for precise analysis of coordinated movement patterns. One disadvantage of the MR is its high cost. The MR apparatus is expensive and has a substantial footprint, requiring a larger dedicated space. Operating the MR is labor-intensive because mice are not inherently motivated to walk across the platform without interruption.

It is important to note that other behavioral paradigms may offer insight into motor function and coordination in an unbiased manner, such as the accelerating Rotarod (Deacon, 2013), which is based on the mouse’s ability to move and balance on an accelerating rod. The time spent on the accelerating road is automatically measured by the apparatus, effectively excluding the operator bias. A drawback of this apparatus is that mice exhibiting significant paralysis, particularly in the early days after SCI, are often incapable of performing the task, rendering this test mostly suitable for chronic injury time points. Moreover, for animals that manage to stay on the rod, the test quickly shifts from a measure of coordination to one of endurance (Deacon, 2013). Alternative assessments, such as the string test, coat hanger test, and horizontal bar test only evaluate forelimb strength and coordination, not locomotion. Additionally, machine-learning algorithms for automated unbiased analyses of these behaviors have been implemented only on a limited scale, with observations specifically focusing on hindlimb behaviors (Eisdorfer et al., 2022; Sato et al., 2022) or grasping behaviors (Duque et al., 2023; O’Neill et al., 2022). Unlike these previous applications, our study comprehensively investigated various unexplored parameters such as forelimb and tail poses, compensatory mechanisms, alterations in head position, lift and stance profiles, anxiety, and fatigue-like behaviors. Moreover, the utilization of the MB addresses inherent motivation issues by gathering relevant motor data without requiring mice to perform specific tasks, such as traversing across a platform.

Ultimately, while the BMS remains a valuable tool to evaluate locomotion in easy-to-execute, cost-effective fashion, its integration with MB and MR will provide a much wider range of information on specific aspects of locomotion that the BMS is unable to extract. Being fully automated, MB and MR are sensitive and devoid of investigator-dependent bias and variability, making them essential tools for ensuring rigor and reproducibility of behavioral outcomes after SCI. In the future, the correlation of MB and MR metrics with molecular, histological, and biochemical analyses at various time points after injury will help establish whether specific MB and/or MR metrics may be predictive of long-term injury outcomes and progression. This will further strengthen the utility of these approaches in ensuring data robustness, which is the foundation on which to plan well-designed clinical studies and trials, thereby increasing the chances of successful outcomes.

In short, deep-learning-based MB and MR tests are powerful tools for understanding disease development and recovery, with extensive applicability across neurological disorders and beyond.

## Author contributions

MR: investigation, methodology, data curation, formal analysis, writing (original draft); EKT, methodology, data curation; EGV, methodology; DJL: conceptualization, methodology, data curation, formal analysis, writing (review and editing), supervision, funding acquisition; RB: conceptualization, methodology, data curation, formal analysis, writing (review and editing), supervision, funding acquisition.

## Data sharing

Our DLC workflow is publicly available in the GitHub repository at https://github.com/MaureenAscona/DLC-MotorBox/tree/main and https://github.com/MaureenAscona/DLC-MotoRater

## Acknowledgments

We thank Jose Mier for his assistance in animal husbandry, Brianna Carney and Antonella Mini for assistance in all perfusions and dissections, Maria Quiala Acosta for administering all spinal cord injuries, and Ramon German and Miguel Martinez for performing all BMS experiments.

