## Supplementary Figures for "A deep learning-based approach for unbiased kinematic analysis in CNS injury"

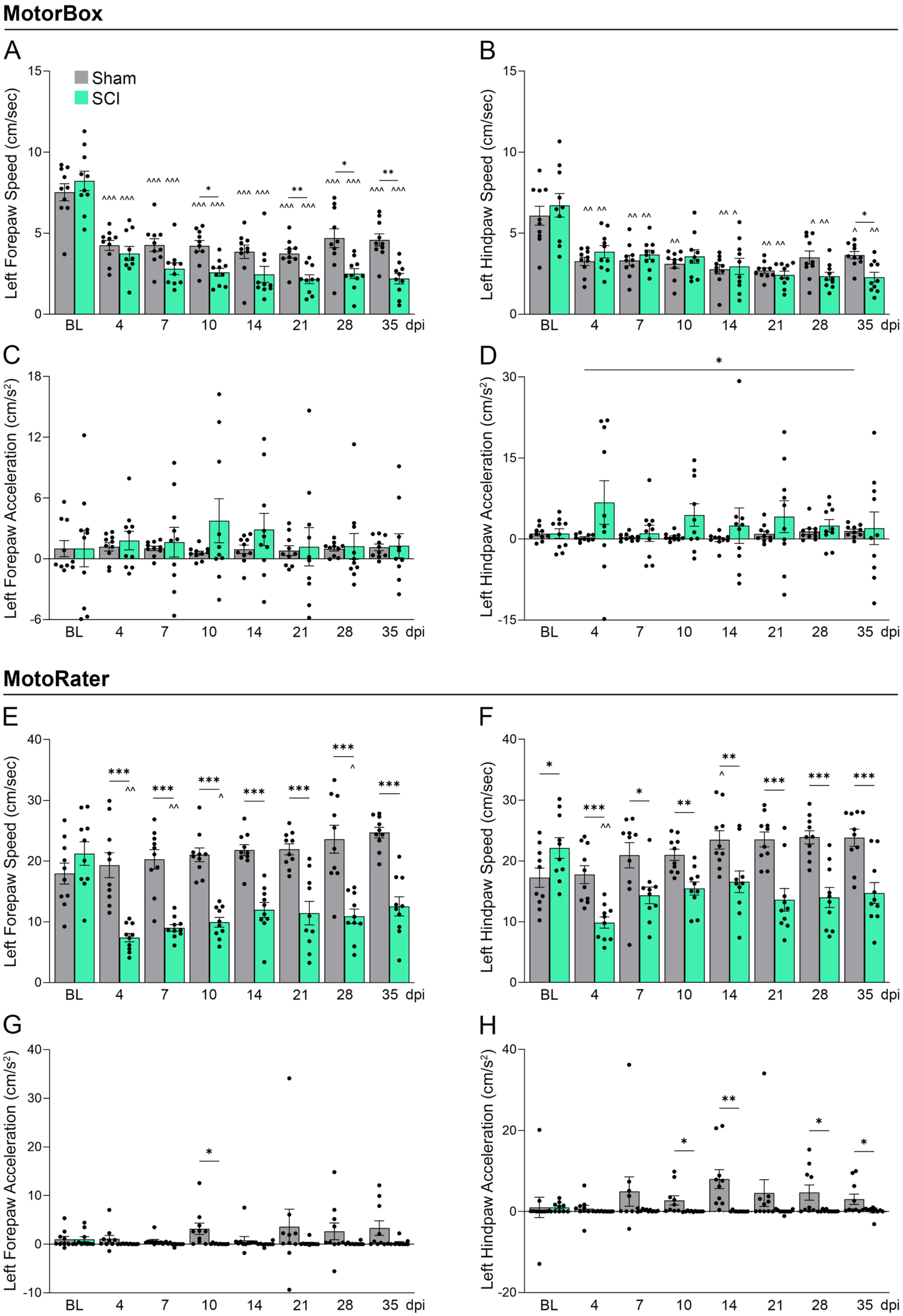

**Supplementary Figure 1. Detection of SCI-induced changes in ambulation metrics for the left side.** (A, B, E, F) Left speed metrics: forepaw speed for (A) MB and (E) MR, hindpaw speed for (B) MB and (F) MR. (C, D, G, H) Left acceleration metrics (calculated using velocity): forepaw acceleration for (C) MB and (G) MR, hindpaw acceleration for (D) MB and (H) MR. Acceleration values were normalized to baseline; n=10/group, *$p \leq 0.05$, **$p \leq 0.01$, ***$p \leq 0.001$ between sham and SCI. ^$p \leq 0.05$, ^^$p \leq 0.01$, ^^^$p \leq 0.001$ compared to BL, Repeated Measures Two-Way ANOVA.

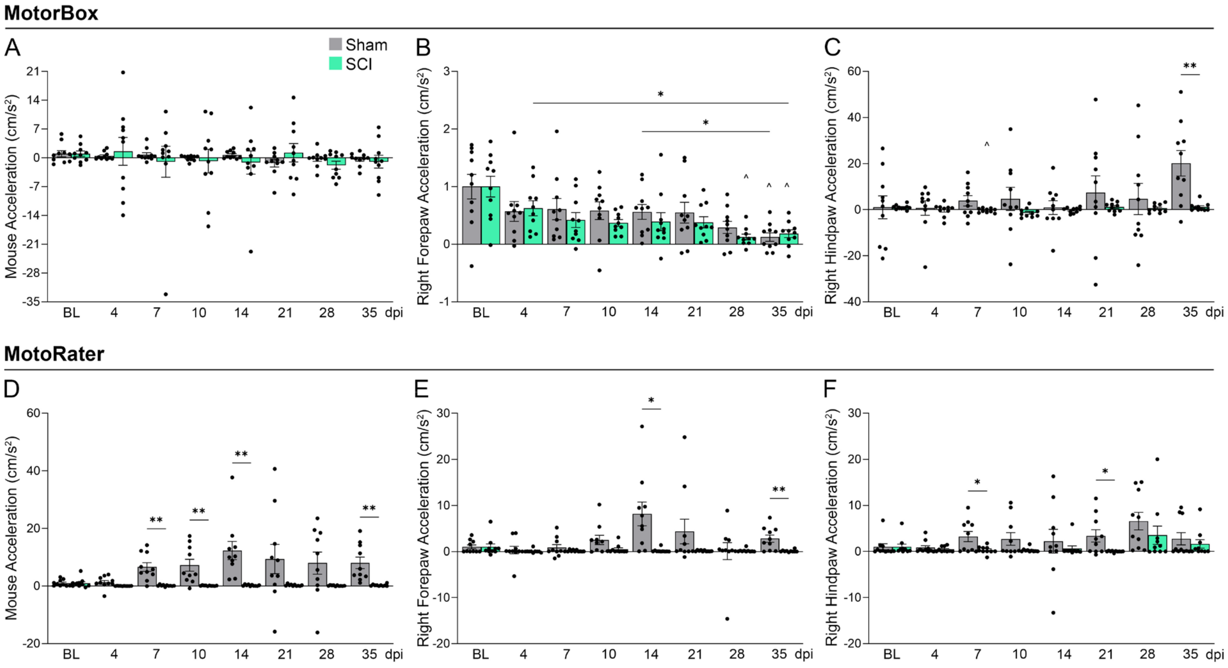


**Supplementary Figure 2. Detection of SCI-induced changes in acceleration metrics for the right side.** (A, D) Mouse acceleration (based on base tail mark) for (A) MB and (D) MR. (B, C, E, F) Right acceleration metrics: forepaw acceleration for (B) MB and (E) MR, hindpaw acceleration for (C) MB and (F) MR. Acceleration values were normalized to baseline; n=10/group, *p≤0.05, **p≤0.01 between sham and SCI. ^p≤0.05 compared to BL, Repeated Measures Two-Way ANOVA

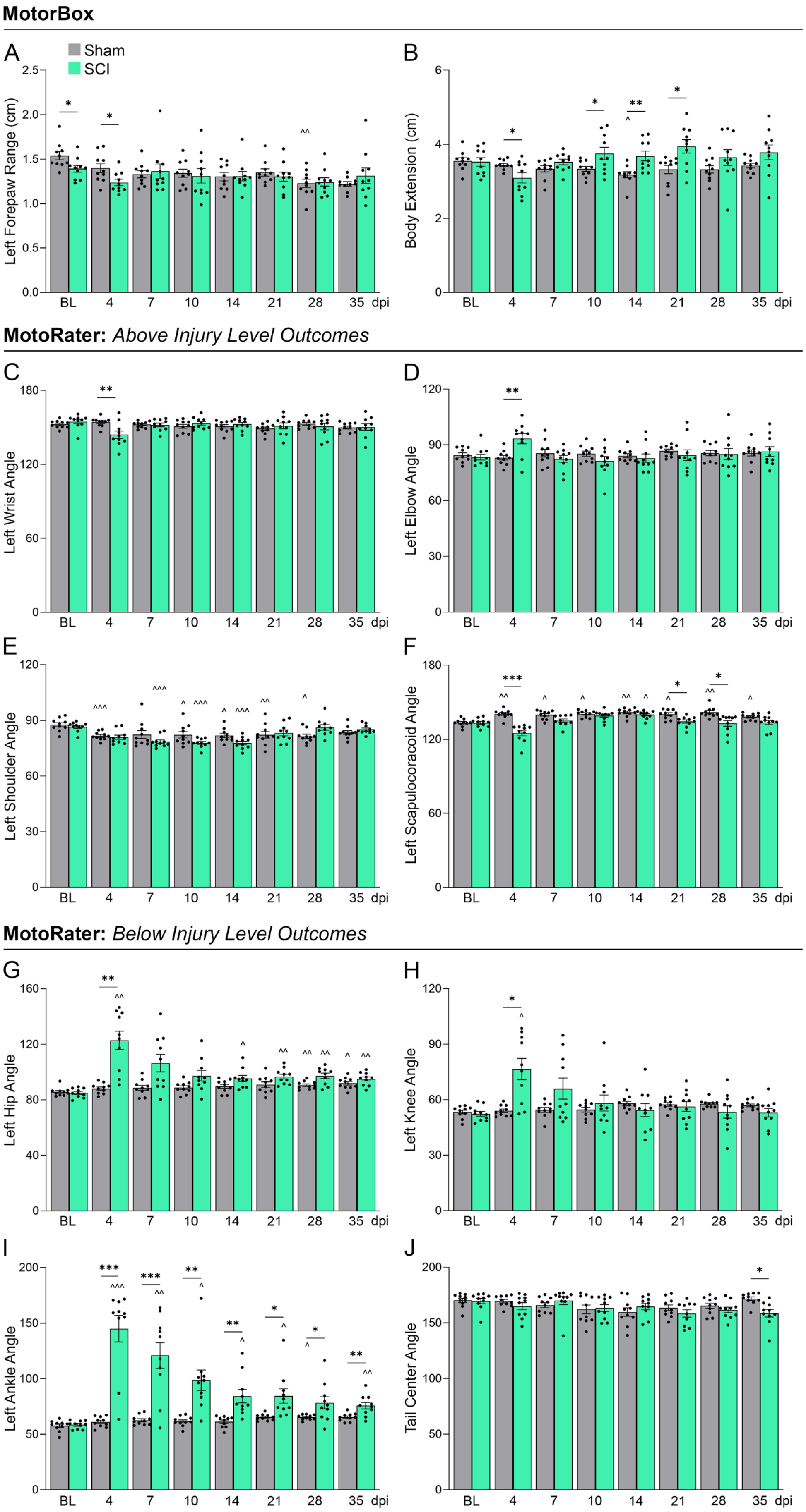

MotorBox
A
Sham
SCI
Left Forepaw Range (cm)
B
Body Extension (cm)
BL
4
7
10
14
21
28
35
dpi
MotoRater: Above Injury Level Outcomes
C
Left Wrist Angle
D
Left Elbow Angle
E
Left Shoulder Angle
F
Left Scapulocoracoid Angle
MotoRater: Below Injury Level Outcomes
G
Left Hip Angle
H
Left Knee Angle
I
Left Ankle Angle
J
Tail Center Angle

**Supplementary Figure 3. Detection of SCI-induced changes in limb range and joint angles.** (A) Forepaw range measured as distance from the left forepaw to the top body in the MB. (B) Body extension measured as distance from the top body mark to the bottom body mark in the MB. (C-F) Angles of above injury level joints in the MR: (C) left wrist, (D) left elbow, (E) left shoulder, and (F) left scapulocoracoid angles. (G-J) Angles of below injury level joints in the MR: (G) left hip, (H) left knee, (I) left ankle, and (J) tail center; n=10/group, *$p \leq 0.05$, **$p \leq 0.01$, ***$p \leq 0.001$ between sham and SCI. ^$p \leq 0.05$, ^^$p \leq 0.01$, ^^^$p \leq 0.001$ compared to BL, Repeated Measures Two-Way ANOVA.

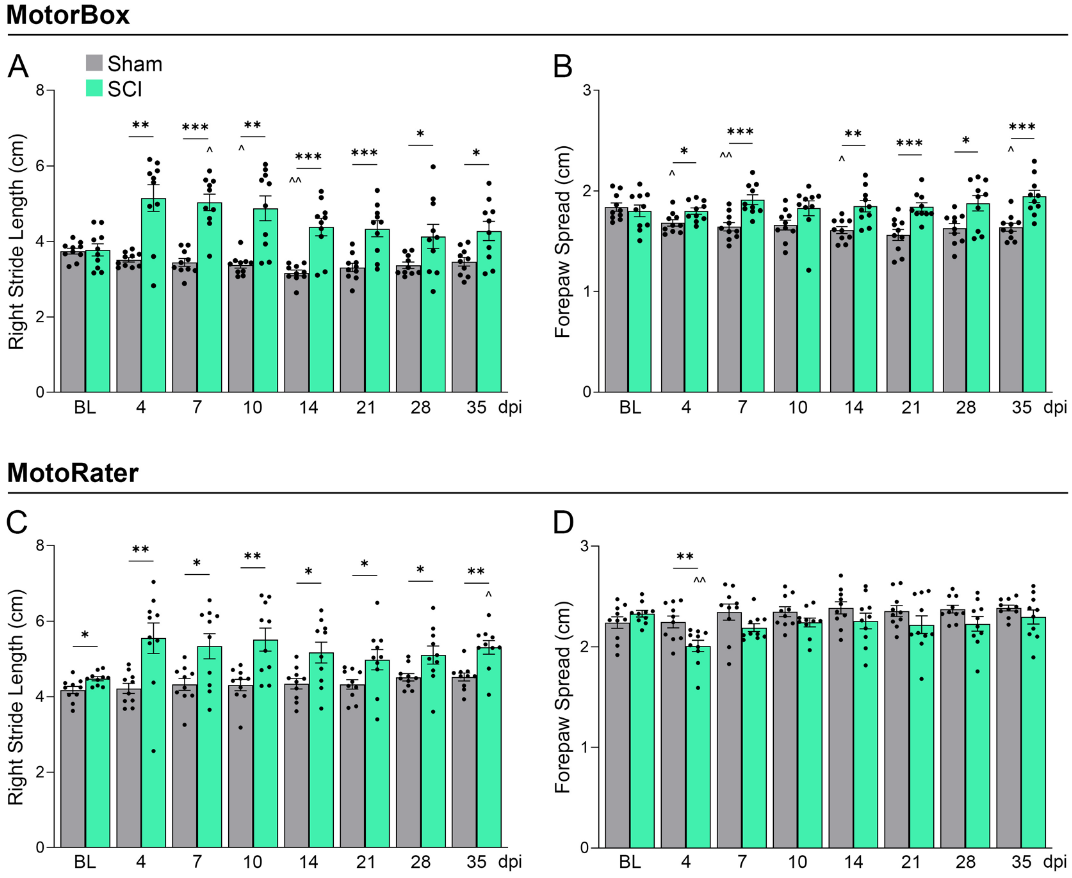


**Supplementary Figure 4. Detection of SCI-induced changes in gait.** (A, C) right stride length measured as distance from the right forepaw to the right hindpaw in the (A) MB, and the (C) MR. (B, D) Forepaw spread measured as the distance from the right forepaw to the left forepaw in (B) MB and (D) MR; n=10/group, *p≤0.05, **p≤0.01, ***p≤0.001 between sham and SCI. ^p≤0.05, ^^p≤0.01 compared to BL, Repeated Measures Two-Way ANOVA.

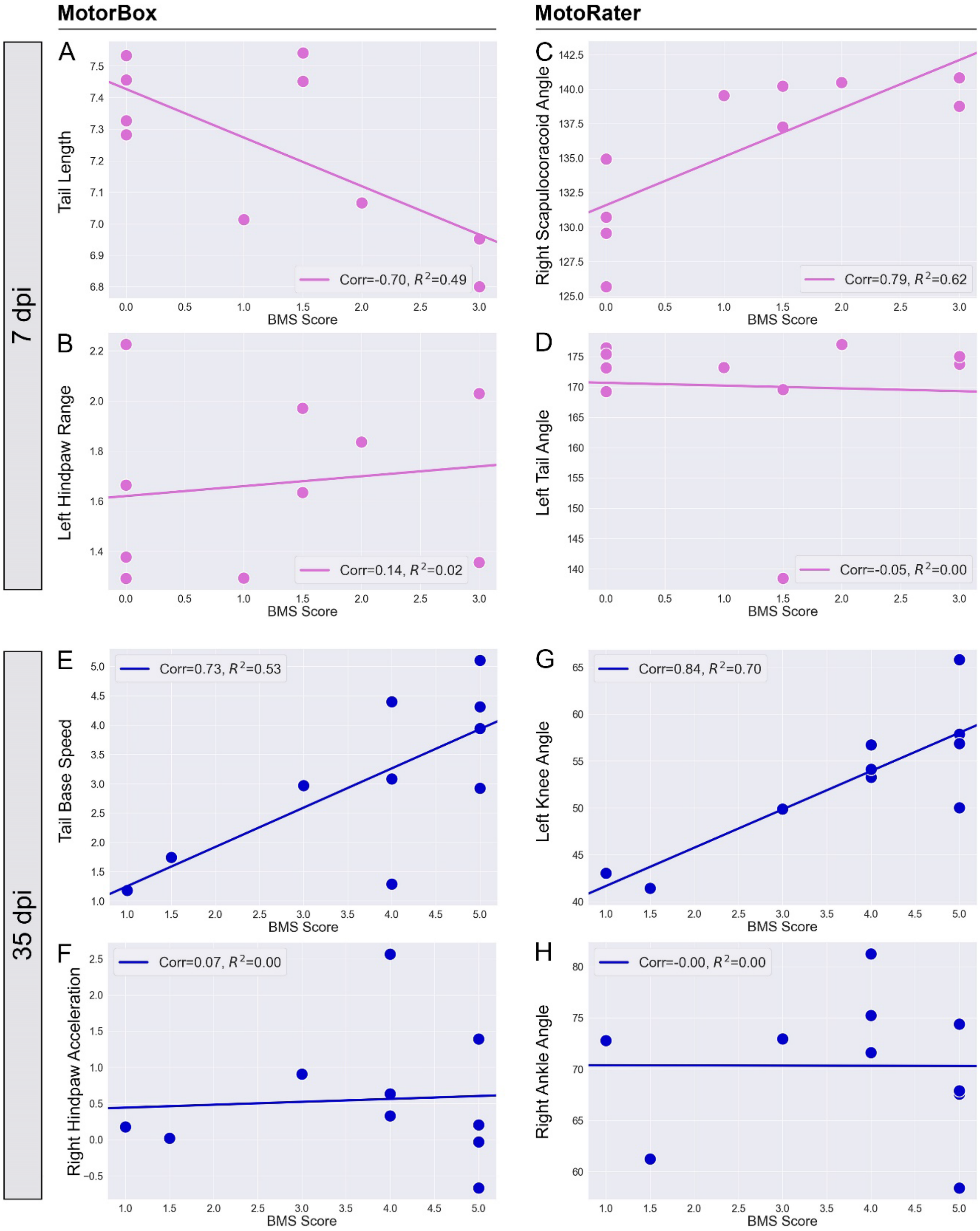


**Supplementary Figure 5. Correlation analysis of select metrics in MB and MR with BMS score at acute and chronic SCI.** (A) Tail length vs BMS score at 7 dpi in MB. (B) Left hindpaw range vs BMS score at 7 dpi in MB. (C) Right scapulocoracoid angle vs BMS score at 7 dpi in

MR. (D) Left tail angle vs BMS score at 7 dpi in MR. (E) Tail base speed vs BMS score at 35 dpi in MB. (F) Right hindpaw acceleration vs BMS score at 35 dpi in MB. (G) Left knee angle vs BMS score at 35 dpi in MR. (H) Right ankle angle vs BMS score at 35 dpi in the MR; n=10/group; Pearson correlations were used to compare MR and MB metrics against BMS scores, while linear regression analyses were used to calculate $R^2$ values.
